# Lipid transport by *Candida albicans* Dnf2 is required for hyphal growth and virulence

**DOI:** 10.1101/2022.06.10.495726

**Authors:** Bhawik K. Jain, Andrew S. Wagner, Todd B. Reynolds, Todd R. Graham

**Author notes:** To whom correspondence should be addressed: Todd R. Graham: Department of Biological Sciences, Vanderbilt University Nashville, TN 37235.

## Abstract

*Candida albicans* is a common cause of human mucosal yeast infections, and invasive candidiasis can be fatal. Antifungal medications are limited, but those targeting the pathogen cell wall or plasma membrane have been effective. Therefore, virulence factors controlling membrane biogenesis are potential targets for drug development. P4-ATPases contribute to membrane biogenesis by selecting and transporting specific lipids from the extracellular leaflet to the cytoplasmic leaflet of the bilayer to generate lipid asymmetry. A subset of heterodimeric P4-ATPases, including Dnf1-Lem3 and Dnf2-Lem3 from *Saccharomyces cerevisiae*, transport phosphatidylcholine (PC), phosphatidylethanolamine (PE), and the sphingolipid glucosylceramide (GlcCer). GlcCer is a critical lipid for *Candida albicans* polarized growth and virulence, but the role of GlcCer transporters in virulence has not been explored. Here we show that the *Candida albicans* Dnf2 (*Ca*Dnf2) requires association with *Ca*Lem3 to form a functional transporter and flip fluorescent derivatives of GlcCer, PC and PE across the plasma membrane. Mutation of conserved substrate-selective residues in the membrane domain strongly abrogates GlcCer transport and partially disrupts PC transport by CaDnf2. *Candida* strains harboring *dnf2* null alleles (*dnf2*ΔΔ) or point mutations that disrupt substrate recognition exhibit defects in the yeast to hyphal growth transition, filamentous growth and virulence in systemically infected mice. The influence of *CaDNF1* deletion on the morphological phenotypes is negligible although the *dnf1*ΔΔ *dnf2*ΔΔ strain was less virulent than the *dnf2*ΔΔ strain. These results indicate that the transport of GlcCer and/or PC by plasma membrane P4-ATPases is important for pathogenicity of *Candida albicans*.

## Introduction

*Candida albicans* is the most common opportunistic pathogenic fungi in humans. It causes mucosal infections in oral, esophageal, and vulvovaginal regions, and life-threatening systemic infections (1–3). The virulence of pathogenic fungi depends on the secretory pathway to transport virulence factors to the exoplasmic side of the plasma membrane for biofilm formation, fungal hydrolase secretion, adhesins and invasins expression and transport of antigenic glycosphingolipids (4–6). The *Candida* secretory pathway also helps control the specific protein and lipid composition of membranes required for virulence and morphological transitions from the budding yeast form to the hyphal growth phase that facilitates infection. For example, the sphingolipid glucosylceramide (GlcCer) is associated with pathogenicity in *Candida albicans* as well as *Cryptococcus neoformans* and *Aspergillus fumigatus* (7–9). GlcCer is also critical for growth, yeast-to-hyphae transition, and hyphal elongation in *Candida albicans, Cryptococcus neoformans, Aspergillus nidulans* and in the plant pathogen *Fusarium graminearum* (8–10). Phosphatidylserine (PS) and phosphatidylethanolamine (PE) are also critical lipids in *Candida albicans* pathogenesis and the fungal-specific PS synthase (Cho1) has emerged as a potential drug target as most antifungal drugs target membranes or membrane biosynthesis (11, 12).

An important property of biological membranes is lipid asymmetry, which plays critical roles in signal transduction, cell division, apoptosis, and intracellular and extracellular vesicle formation (13). Phosphatidylcholine (PC) and sphingolipids are typically enriched in the extracellular leaflet, and phosphatidylethanolamine (PE) and phosphatidylserine (PS) are mainly concentrated in the cytosolic leaflet of the plasma membrane in mammalian cells (14, 15). In *Saccharomyces cerevisiae*, PC is enriched in the cytosolic leaflet along with PE and PS while the extracellular leaflet is primarily composed of glycosphingolipids (16). P4-ATPases are lipid flippases that transport lipids from the exoplasmic leaflet of the membrane to the cytoplasmic leaflet to maintain this membrane lipid asymmetry (17, 18). Most P4-ATPases are heterodimeric protein complexes with a catalytic α subunit (P4-ATPase) and accessory β subunit (Cdc50/Lem3/Crf1 in budding yeast). For example, Lem3 forms a heterodimer complex with Dnf1 or Dnf2 that is necessary for their exit from the endoplasmic reticulum, and loss of Lem3 impairs Dnf1,2-dependent lipid transport. Humans express 14 P4-ATPases (ATP8A1-ATP11C; CDC50A-C)) whereas budding yeast express 5 P4-ATPases: Dnf1-Lem3, Dnf2-Lem3, Dnf3-Crf1, Drs2-Cdc50 and Neo1, which lacks a β-subunit (13, 17, 19–21).

Mutations in P4-ATPase genes are associated with a number of diseases and the fungal genes are becoming linked to pathogenesis. Human P4-ATPase mutations are linked to neurological disease (ATP8A2, ATP9A, ATP8B2, ATP10B, ATP11A), metabolic and cardiovascular disease (ATP10A, ATP10D), cholestasis and hearing loss (ATP8B1), systemic sclerosis (ATP8B4), severe Covid-19 (ATP11A), cancer progression (ATP11B) and hemolytic anemia (ATP11C)(22, 23, 32–37, 24–31). The P4-ATPase Apt1-Cdc50 from *Cryptococcus neoformans* transports a variety of substrates, including GlcCer, and is essential in polysaccharide secretion, stress tolerance, fungal survival inside macrophages, and virulence (38–41). A *Candida albicans drs2*ΔΔ flippase mutant shows defects in hyphal growth and hypersensitivity to fluconazole (42). Also, *Ca*Cdc50 plays a vital role in antifungal drug resistance, cell membrane functions, hyphal development and virulence (43). The *Aspergillus nidulans* P4-ATPases DnfA and DnfD play roles in hyphal growth (44). In *Magnaporthe oryzae*, the flippases MoPde1 and MoApt2 are involved in fungal virulence (45, 46) and the *F. graminearum* FgDnfA is crucial for vegetative growth and pathogenesis (47). However, the substrate specificity of most of these fungal P4-ATPases has not been defined.

P4-ATPases differ in their substrate specificity and cellular localizations but have similar mechanisms for recognition and transport of lipids. Dnf1 and Dnf2 localize to the plasma membrane and endosomes and transport PC, PE, and the sphingolipid glucosylceramide (GlcCer)(48–51). Drs2, Dnf3 and Neo1 localize primarily to the Golgi apparatus and transport PS and PE (52–56). Like other P-type ATPases, P4-ATPases bind ATP at their cytosolic nucleotide binding (N) domain, transfer phosphate to a conserved aspartic acid residue in the phosphorylation (P) domain, and subsequently dephosphorylate themselves using residues in the actuator (A) domain. This ATP-dependent reaction cycle induces conformational changes in the membrane (M) domain, particularly transmembrane (TM) segments 1-4, that drives unidirectional lipid transport(57–59). Substrate lipid is drawn from the exoplasmic leaflet into an “entry gate” binding site formed by specific substrate-selective TM residues identified through mutational approaches (21, 60–64). The substrate is then flipped and docked at an “exit gate” site before release to the cytosolic leaflet (17, 18, 65). Recent cryo-EM structures of several P4-ATPases revealed the lipid positions in the two gates and identified additional amino acids important for substrate selection (55, 57–59, 66, 67). GlcCer transporter activity was first discovered for Dnf1 and Dnf2 from *S. cerevisiae* and is a conserved function of human P4-ATPases in the ATP10 subgroup (48). However, *S. cerevisiae* lacks the GlcCer synthase enzyme and does not synthesize this sphingolipid (68). Therefore, the role of GlcCer transport in fungal cell physiology is unclear.

*Candida albicans* expresses GlcCer synthase (Hsx11)(8, 68) and five putative P4-ATPases, two of which appear to be orthologous to the Dnf1-Lem3 and Dnf2-Lem3 PC/GlcCer transporters in *Saccharomyces cerevisiae*. In this study, we functionally characterized the *Candida albicans* Dnf1-Lem3 and Dnf2-Lem3 and our data indicate that these P4-ATPases transport PC, PE and GlcCer comparably to their *S. cerevisiae* orthologs. We also report that *Ca*Dnf2 makes an important contribution to hyphal growth and virulence.

## Results

### The *Candida albicans* genome encodes 5 P4-ATPases

A Protein BLAST analysis of the *Candia albicans* proteome queried using P4-ATPases from budding yeast indicates that *Candida albicans* encodes 5 P4-ATPases. Based on sequence homology, phylogenetic analysis, and functional studies described below, we named the *Candida albicans* P4-ATPase α subunits as *Ca*Dnf1 (C3_03250W_A), *Ca*Dnf2 (C5_00570W_A), *Ca*Dnf3 (C4_03110W_A), *Ca*Drs2 (C3_07230W_A), and *Ca*Neo1 (C1_04630C_A)(Fig1A, FigS1). Prior phylogenetic studies suggested that P4-ATPase alpha subunits can be classified into two clades, the P4A-ATPases containing the fungal Dnf and Drs2 proteins, and the P4B-ATPases containing fungal Neo1 proteins (69). The phylogenetic analysis of the accessory β subunits suggests that the *C. albicans* proteins Cdc50 (C6_03660C_A), Lem3 (C2_05110W_A), Crf1 (CR_07600W_A) are orthologous to ScCdc50, ScLem3 and ScCrf1, respectively (Fig1B).

**Figure 1:**
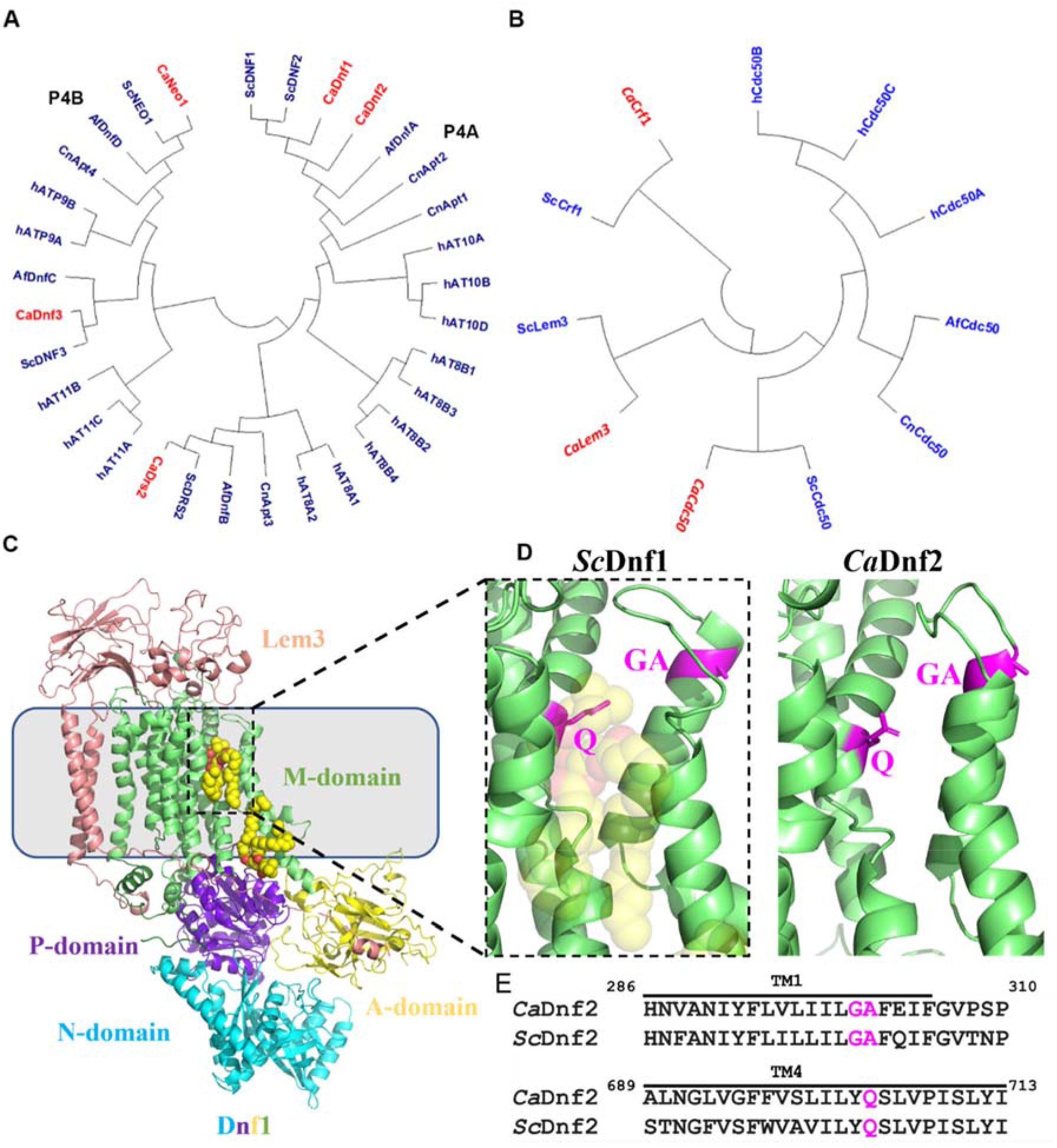
*Candida albicans* potentially expresses 5 P4-ATPases. Phylogenetic tree of P4-ATPase α subunit (A) and β subunit (B) sequences from *Saccharomyces cerevisiae, Candida albicans, Aspergillus fumigatus, Cryptococcus neoformans and Homo sapiens*. The red color indicates the *Candida albicans* P4-ATPases. (C) The structure of *S.cerevisiae* Dnf1-Lem3 with the major domains labeled in different colors (PDB, 7KYC) (58). (D) Enlarged view of substrate translocation path entry gate of *Sc*Dnf1 with a bound PC and that of *Ca*Dnf2 predicted using AlphaFold2. (E) Sequence alignment of TM1 and TM4 region of *Ca*Dnf2 and *Sc*Dnf2. The TM4 Gln and TM1 GlyAla residues in magenta are important for PC/GlcCer transport. Numbers are for the CaDnf2 amino acid sequences.

For P4-ATPases, lipid substrate selection and transport are mediated by specific residues in the M domain. The structure of ScDnf1-Lem3 with PC substrate molecules bound in the substrate translocation path and an enlargement of the entry gate near the extracellular side of the membrane is shown in Figure 1C and 1D (58). We aligned the sequence of TM1 and TM4 of *Sc*Dnf2 and *Ca*Dnf2, which showed that *Ca*Dnf2 has the conserved TM1 GA (amino acid 300 and 301) and TM4 Q (amino acid 704) residues known to be important for PC and GlcCer transport in the budding yeast ortholog (Fig. 1E). The small side chains of the TM1 GA appear to provide a permissive environment for PC and GlcCer binding whereas the TM4 Gln is particularly crucial for GlcCer recognition (21, 48, 58, 60). To further understand the structure and function of *Ca*Dnf1 and *Ca*Dnf2, we generated a model of the *Ca*Dnf2 structure using AlphaFold2 (70). The predicted structure of the *Ca*Dnf2 entry gate is very similar to that of *Sc*Dnf1 and *Sc*Dnf2 (Fig 1D). In addition, the structural model suggested that the overall architecture of *Ca*Dnf2 is similar to all P4A-ATPases for which structures are known. These analyses suggested that *Ca*Dnf2 would have a transport substrate specificity comparable to *Sc*Dnf1 and *Sc*Dnf2.

### Heterologous expression of *Candida albicans* Dnf2-Lem3 in *S. cerevisiae*

To test if *Ca*Dnf2 can catalyze flippase activity and to define its substrate specificity, we cloned the *CaDNF2* gene under control of the *ScDNF1* promoter for heterologous expression in *S. cerevisiae. Ca*Dnf2 was expressed in a strain harboring deletion of *ScDNF1* and *ScDNF2* (*Scdnf1,2*Δ) but expressing endogenous *Sc*Lem3. Interestingly, we observed that cells expressing *Ca*Dnf2 alone failed to take up NBD-PC or NBD-GlcCer compared to wild-type (WT) cells or the *Scdnf1,2*Δ cells expressing ScDnf2, and were similar to the empty vector (EV) control (Fig2A,B). This result suggested that *Ca*Dnf2 could not form a heterodimeric complex with *Sc*Lem3 to produce a functional transporter. Therefore, we co-expressed *Ca*Dnf2 and *Ca*Lem3 in *S. cerevisiae* and found that the cells displayed a robust NBD-PC and NBD-GlcCer transport activity. The magnitude of *Ca*Dnf2-*Ca*Lem3 catalyzed lipid transport was at the same level catalyzed by *Sc*Dnf2-*Sc*Lem3 suggesting that *Ca*Dnf2 can function similarly to *Sc*Dnf2 (Fig2A,B) in the ability to localize to the plasma membrane and transport both a sphingolipid and a glycerophospholipid.

**Figure 2:**
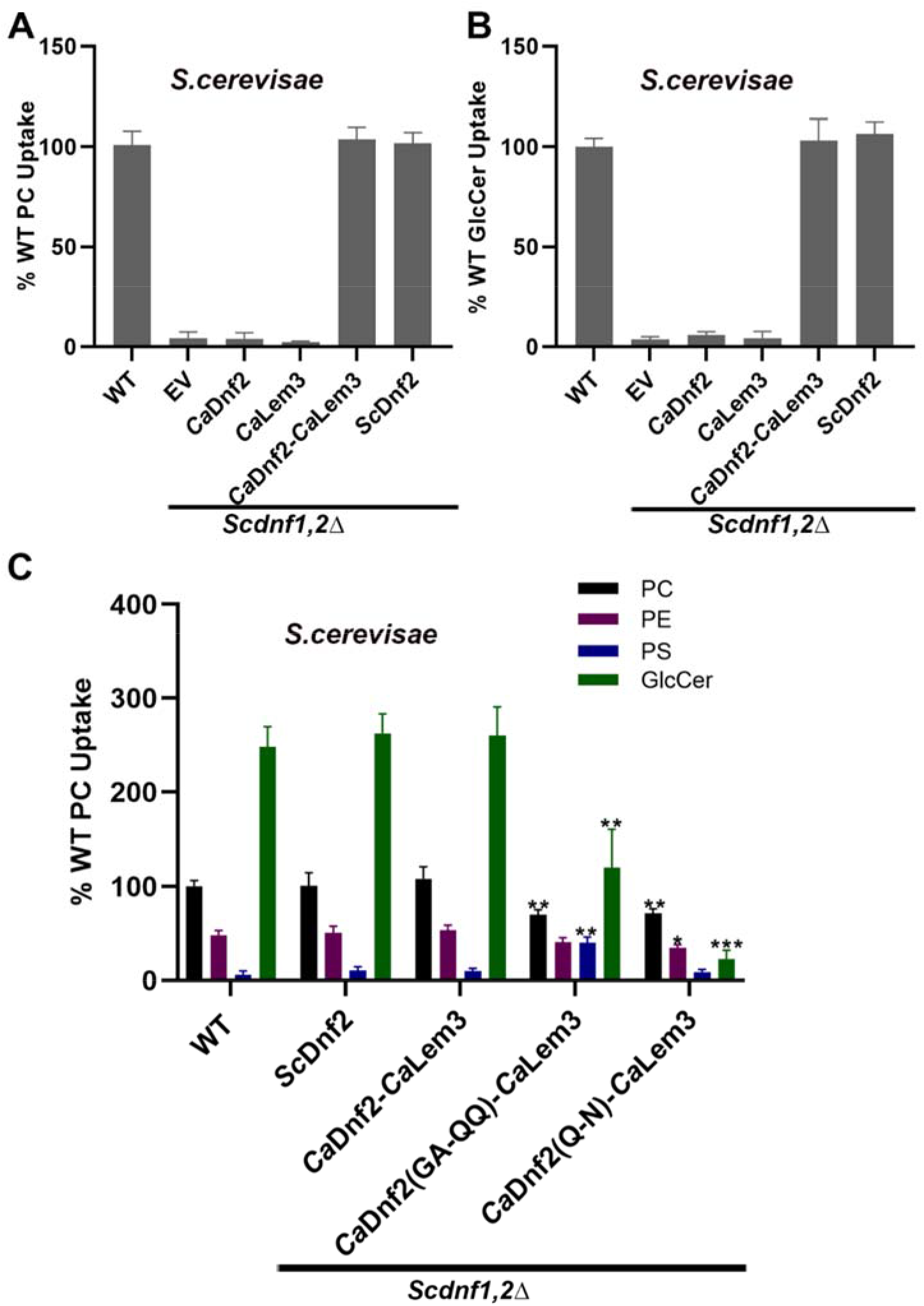
*Ca*Dnf2-*Ca*Lem3 transports GlcCer, PC and PE. NBD-PC (A) and NBD-GlcCer (B) uptake was measured in WT (BY4741) and *dnf1,2*Δ cells expressing empty vector (EV = pRS313), pRS313-*Ca*DNF2, pRS315-*CaLem3* or pRS313-*Sc*DNF2. (C) NBD-PC, NBD-PE, NBD-PS, and NBD-GlcCer uptake was measured in CaDnf2(GA-QQ) and CaDnf2(Q-N) expressed in *Scdnf1,2*Δ. Variance was assessed among data sets using one-way ANOVAs, and comparisons with WT were made with Tukey’s post hoc analysis. *n* ≥ 9, ± S.D *p<0.05, **p<0.01, ***p<0.0005.

We further tested the substrate specificity of *Ca*Dnf2-CaLem3 by measuring its ability to catalyze uptake of NBD-PC, NBD-PE, NBD-PS and NBD-GlcCer. Transport of each substrate was normalized to the amount of NBD-PC taken up by WT *Saccharomyces cerevisiae* and compared to *dnf1*Δ *dnf2*Δ cells expressing *Sc*Dnf2-*Sc*Lem3. Almost identically to its *Saccharomyces cerevisiae* ortholog, *Ca*Dnf2-*Ca*Lem3 mediated uptake of NBD-GlcCer at a rate 2.5-fold greater than NBD-PC and 5-fold greater than NBD-PE. NBD-PS uptake was not significantly different than background for either WT transporter. Thus, *Ca*Dnf2-Lem3 is a GlcCer/PC/PE transporter (Fig2C).

Mutation of the *Sc*Dnf2 TM1 GA or TM4 Q to residues present in PS flippases (TM1 GA to QQ and TM4 Q to N), modestly abrogates PC transport and strongly perturbs GlcCer transport (48). We mutated these residues in *Ca*Dnf2 and observed that *Ca*Dnf2(GA-QQ) and *Ca*Dnf2(Q-N) mutants similarly showed reduced transport of PC and GlcCer. The *Ca*Dnf2(GA-QQ) mutant displayed a 50% reduction in PC transport and 60% reduction in GlcCer transport activity while *Ca*Dnf2(Q-N) mutants showed an approximately 60% reduction in PC transport and 90% reduction in GlcCer transport (Fig2C). As previously observed for ScDnf1-Lem3, the CaDnf2 GA-QQ mutation in TM1 significantly increased the transport of NBD-PS without changing NBD-PE transport. These results indicate that *Ca*Dnf2-Lem3 can transport PC and GlcCer using a conserved mechanism for substrate binding and transport.

### Dnf2 primarily catalyzes plasma membrane GlcCer/PC flippase activity in *Candida albicans*

Next, we examined the NBD-PC and NBD-GlcCer transporter activity in *Candida albicans*. A Wild-type strain revealed a robust flippase activity for NBD-PC and NBD-GlcCer (Fig. 3A,B). Deletion of both alleles of *Cadnf1* (*Cadnf1*ΔΔ) or *Cadnf2* (*Cadnf2*ΔΔ) elicited a significant reduction of PC and GlcCer uptake. Interestingly *Cadnf2*ΔΔ showed a more substantial decrease in NBD-PC and NBD-GlcCer uptake compared to *Cadnf1*ΔΔ indicating *Ca*Dnf2 is the major PC and GlcCer transporter at the plasma membrane. Similarly, ScDnf2 catalyzes the majority of NBD-PC and NBD-GlcCer transport at the plasma membrane of budding yeast. We also created a *Cadnf1*ΔΔ *Cadnf2*ΔΔ (*Cadnf1,2*ΔΔ) strain, which showed a stronger reduction in PC and GlcCer transport activity. NBD-lipid transport activity was restored upon expression of *CaDNF2* (Fig.3A,B). These data indicate that *Candida albicans* Dnf2 transports PC and GlcCer in the native membrane environment comparably to what was observed when expressed in *Saccharomyces cerevisiae*. We also expressed *Ca*Dnf2(GA-QQ) and *Ca*Dnf2(Q-N) mutants in the *Cadnf1,2*ΔΔ strain and again observed that the TM1 GA motif and conserved TM4 glutamine is critical for PC and GlcCer transport activity (Fig.3C,D). The influence of the mutations on PC transport was more pronounced when assayed in *Candida albicans* than when the same *Ca*Dnf2 mutant variants were assayed in *Saccharomyces cerevisiae*. We suspect that this reflects an influence of the different membrane compositions on transporter activity.

**Figure 3:**
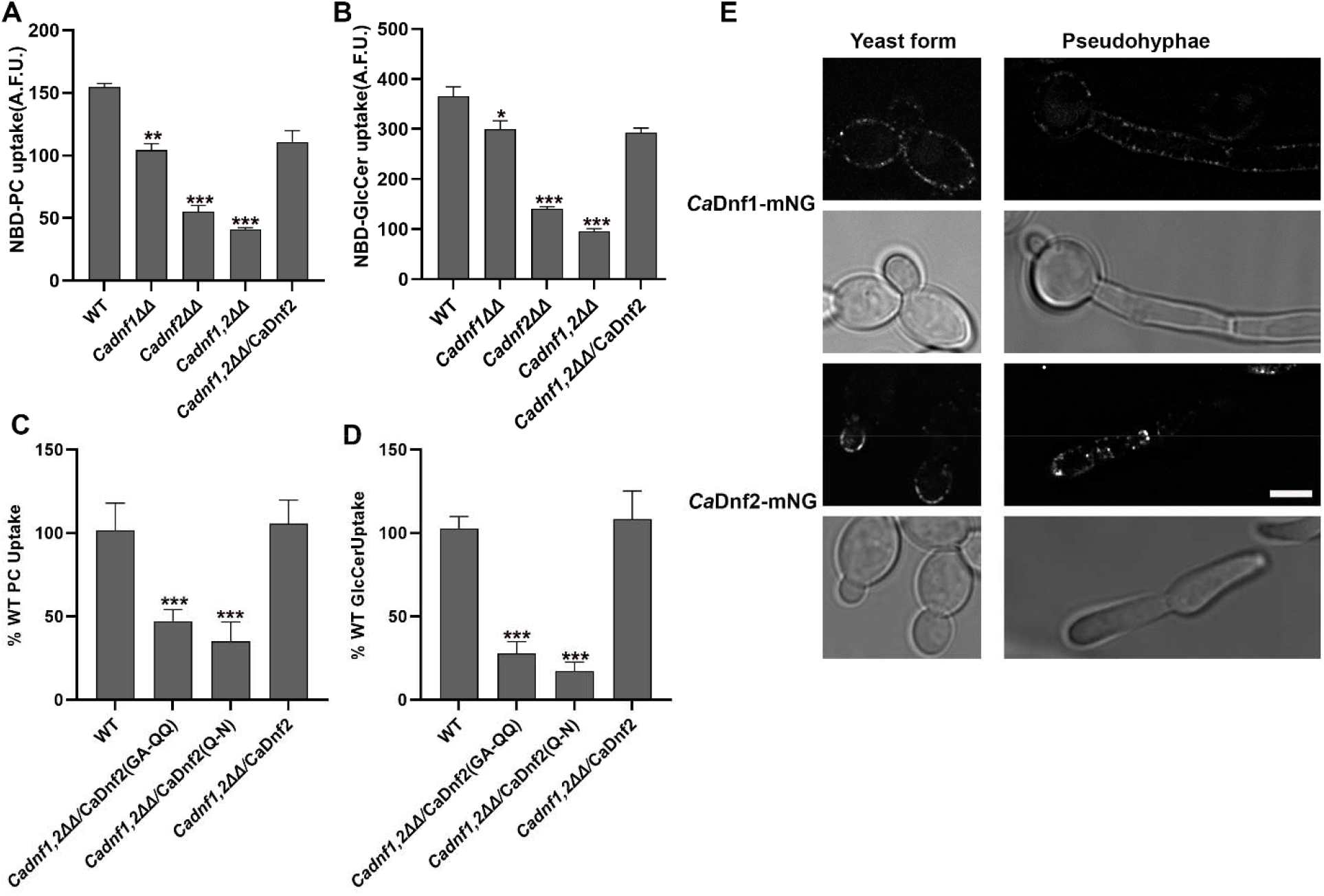
Dnf1/2 transports PC and GlcCer across the *Candida albicans* plasma membrane. NBD-PC (A) and NBD-GlcCer uptake was measured in WT, *Cadnf1*ΔΔ, *Cadnf2*ΔΔ, *Cadnf1,2*ΔΔ and *Cadnf1,2*ΔΔ cells expressing *Ca*DNF2 and presented as raw arbitrary fluorescent units (A.F.U.) without background subtraction or normalization (n ≥ 9) ± S.D. (error bars)*. Ca*DNF2, *Ca*DNF2(GA-QQ) or *Ca*DNF2(Q-N) was reintroduced in the *Cadnf1,2*ΔΔ strains and NBD-PC (C) and NBD-GlcCer (D) uptake was measured. Fluorescence uptake in the *Cadnf1,2*ΔΔ strain was subtracted as background and data were normalized to the WT strain. Variance was assessed among data sets using one-way ANOVAs, and comparisons with WT were made with Tukey’s post hoc analysis. *n* ≥ 9, ± S.D *p<0.05, **p<0.01, ***p<0.0005. (E) Localization of *Ca*Dnf1-mNG and *Ca*Dnf2-mNG in yeast form and hyphal form. Scale bar = 2 μm.

To examine the localization of *Ca*Dnf1 and *Ca*Dnf2 in *Candida albicans*, we tagged *Ca*Dnf1 and *Ca*Dnf2 with mNeonGreen. We observed that *Ca*Dnf1 and *Ca*Dnf2 mostly localized to the plasma membrane in the yeast form and remained primarily at the plasma membrane in hyphal cells upon induction of filamentous growth. Whereas *Ca*Dnf1-mNG was uniformly distributed around mother and daughter cells, *Ca*Dnf2 displayed a polarized localization to the daughter cell of both yeast form and hyphal cells (Fig.3E).

### *Candida albicans* Dnf2 flippase activity contributes to hyphal growth

P4-ATPases have been associated with polarized growth (50, 51, 56, 71); therefore, we examined the role of *Ca*Dnf1 and *Ca*Dnf2 in hyphal growth by growing the mutant strains on spider medium at 37°C for 7 days (Fig. 4). On spider medium, the hyphal growth for *Cadnf2*ΔΔ was significantly reduced compared to the WT strain. In contrast, *Cadnf1*ΔΔ did not show a defect in hyphal growth or colony morphology. The *Cadnf1,2*ΔΔ strain also showed a substantial reduction in colony size but was similar to the *Cadnf2*ΔΔ strain. The hyphal growth phenotype was restored to wild type levels by introducing *CaDNF2* in the *Cadnf1,2*ΔΔ strain (Fig.4A). These results suggest that *Ca*Dnf2 is the major plasma membrane flippase contributing to hyphal growth. Interestingly, when we introduced *Ca*Dnf2(GA-QQ) and *Ca*Dnf2(Q-N) mutants with reduced PC and GlcCer transport activity in *Cadnf1,2*ΔΔ strains they failed to restore the hyphal growth phenotype (Fig4A). The colony size was also significantly reduced in *Cadnf2*ΔΔ strain and PC and GlcCer transport deficient mutants (Fig 4C). The colony size of the *Cadnf1*ΔΔ strain was not significantly different from WT, and the *Cadnf1,2*ΔΔ double mutant was no different from the *Cadnf2*ΔΔ single mutant. These data suggest that flippase activity of *Ca*Dnf2 is important for hyphal growth while *Ca*Dnf1 is dispensable.

**Figure 4:**
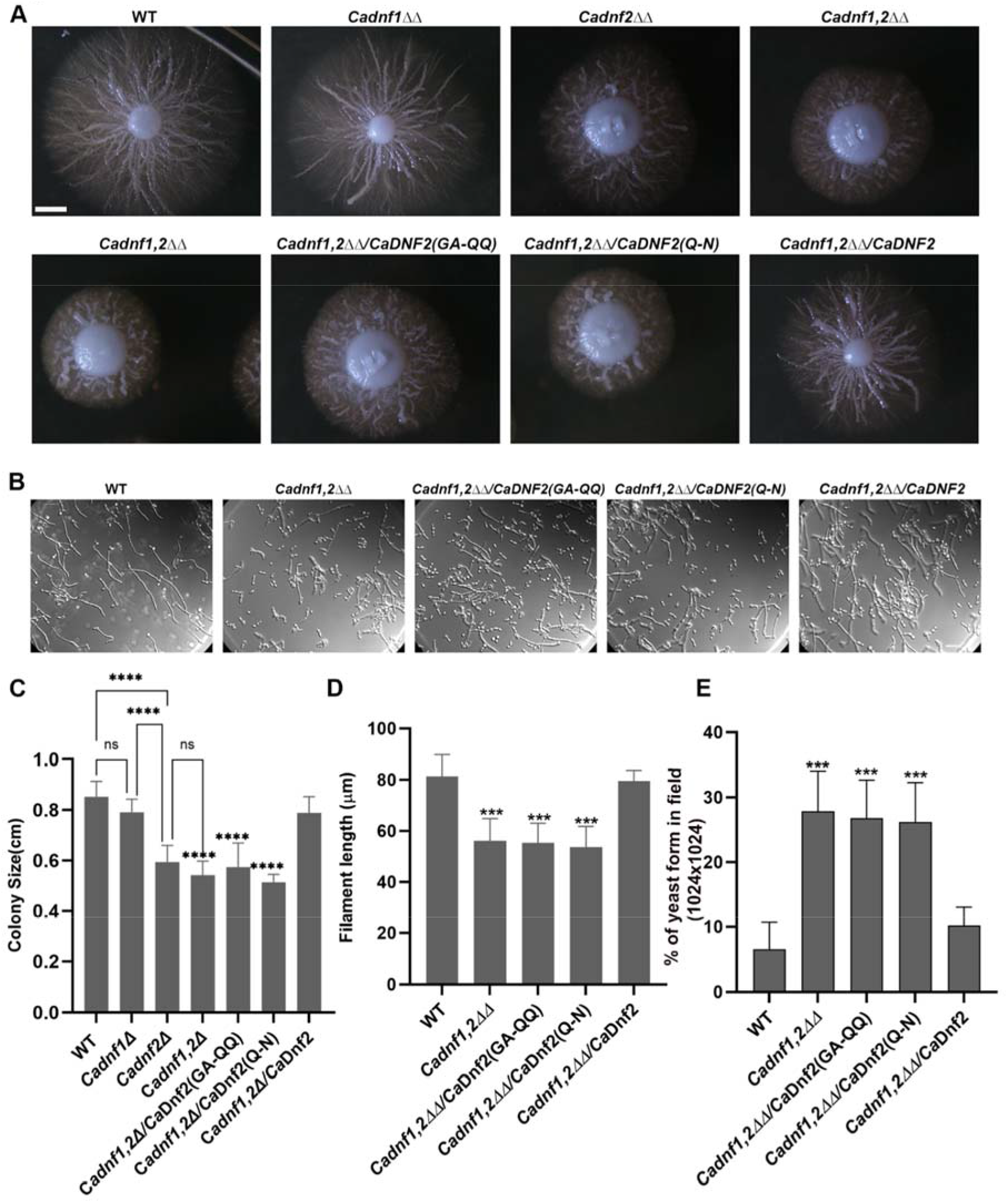
*Candida albicans* Dnf1/2 flippase activity is important for dimorphic growth. **(A)** Filamentation of WT strain and *Cadnf1*ΔΔ, *Cadnf2*ΔΔ and *Cadnf1,2*ΔΔ strains on solid spider media. The images were captured after 7 days incubation at 37°C. Scale bar = 2 mm (B) Filamentous growth in the liquid media after inducing the filamentation for 4 hrs at 37°C in Wild type and mutants. The colony size (C) and filament length was reduced in P4-ATPase mutant strains. Scale bar = 10μm. (E) Number of yeast forms was increased in P4-ATPases mutant compared to WT strain. Variance was assessed among data sets using one-way ANOVAs, and comparisons with WT were made with Tukey’s post hoc analysis. *n* ≥ 30, ± S.D *p<0.05, **p<0.01, ***p<0.0005.

To further understand the role of *Ca*Dnf1/2 in hyphal development, we also examined filamentous growth in liquid spider media by inducing the filamentation at 37°C for 4 hr. The wild-type strains formed regular hyphae upon filamentous growth induction. However, the *Cadnf1,2*ΔΔ mutant showed a strong defect in hyphae formation (Fig. 4B). The hyphal filament length was significantly reduced in the *Cadnf1,2*ΔΔ mutant compared to the wild-type strain (Fig. 4D). The introduction of *Ca*DNF2 in the *Cadnf1,2*ΔΔ strain restored the normal filamentation but the flippase activity mutants failed to restore the filamentation. Hyphal filament length was also reduced in *Ca*Dnf2(GA-QQ) and *Ca*Dnf2(Q-N) strains compared to the WT strain (Fig. 4B,D). We also counted the percentage of cells in the yeast form after induction of filamentation. *Cadnf1,2ΔΔ, Ca*Dnf2(GA-QQ) and *Ca*Dnf2(Q-N) showed a significant increase in yeast form cells in comparison to the WT strain (Fig. 4E). Our data suggest that *Ca*Dnf2 plays a critical role in the transition from yeast form to hyphal growth and its flippase activity is important for hyphal elongation.

### *CaDnf2* flippase activity is important for virulence

To test the role of *Ca*Dnf1/2 in *Candida albicans* virulence, we intravenously infected mice with wild type (WT), *Cadnf1ΔΔ, Cadnf2*ΔΔ, and *Cadnf1,2*ΔΔ strains. Mice were sacrificed 5 days post-infection and we observed a significant reduction in fungal burden in the kidneys of mice harboring the *Cadnf2*ΔΔ and *Cadnf1,2*ΔΔ mutant strains compared to WT strains (Fig5A). Mice injected with the *Cadnf1,2*ΔΔ strain where *Ca*DNF2 was reintroduced showed similar fungal burden as mice injected with the WT strain. In contrast, mice harboring the flippase activity deficient mutants, *Ca*Dnf2(GA-QQ) or *Ca*Dnf2(Q-N), showed a significant decrease in fungal burden and failed to phenocopy the wild-type strain (Fig 5B). These data suggest that *Ca*Dnf1/2 flippase activity is essential for *Candida albicans* virulence.

**Figure 5:**
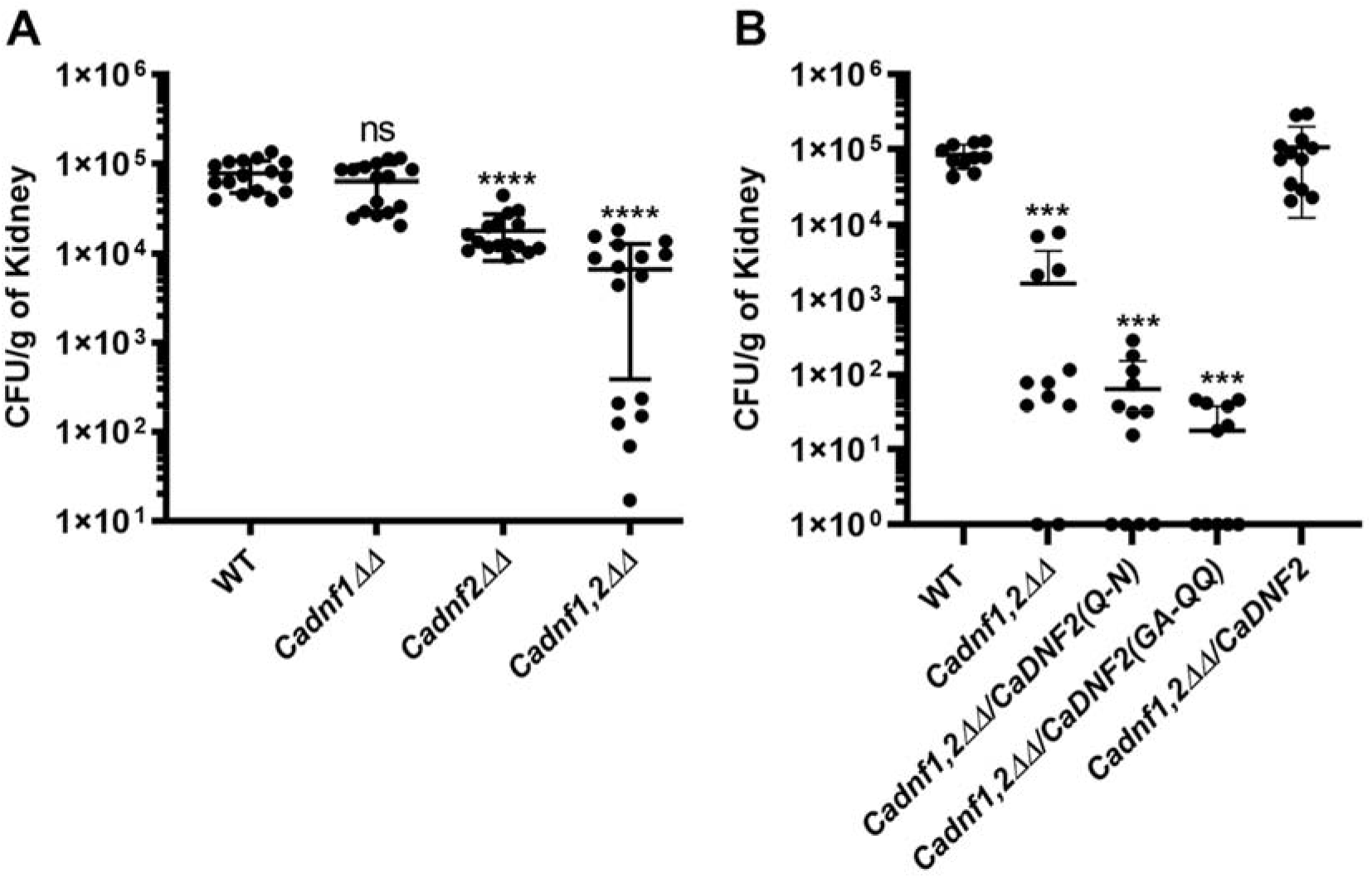
*Ca*Dnf1/2 flippase activity is essential for *Candida albicans* virulence. (A) ICR mice were intravenously infected with 1×10^6^ cells of *C*. *albicans* WT or *Cadnf1*ΔΔ, *Cadnf2*ΔΔ and *Cadnf1,2*ΔΔ strains. The kidneys were harvested 5 days post infection (d.p.i.) to assess fungal burden. (n = 8 mice). (B) ICR mice were intravenously infected with 1×10^6^ cells of *Cadnf1,2*ΔΔ strains reintroduced with CaDNF2, CaDNF2(GA-QQ) or CaDNF2(Q-N) and kidney burden were measured at 5 days post infection. (n = 5-6 mice per strain). ***p<0.0005, ****p<0.0001, by One-way Anova and Holm-Šídák’s multiple comparisons test.

## Discussion

We report here the identification and functional characterization of the *Candida albicans* flippases Dnf1 and Dnf2, which localize to the plasma membrane and transport sphingolipid and glycerophospholipid substrates. The heterologous expression of *Ca*Dnf2-*Ca*Lem3 in *S. cerevisiae* suggests that these proteins must be co-expressed to form a functional flippase that transports, in order of substrate preference, GlcCer, PC and PE. Mutation of conserved substrate selection residues in TM1 and TM4 of *Ca*Dnf2 alter the substrate preference of the flippase and strongly ablate GlcCer transport. Deletion of *Ca*Dnf2 or mutation of residues mediating substrate selection reduces hyphal growth. Furthermore, virulence was attenuated in *Candida albicans* strains lacking *Ca*Dnf2 or expressing variants deficient in GlcCer and PC transport.

The phylogenetic, structural, and functional analyses described herein indicate that the *Candida albicans DNF1*/C3_03250W_A and *DNF2*/C5_00570W_A paralogs are orthologous to *S. cerevisiae DNF1* and *DNF2*, respectively. Interestingly, it was difficult to ascertain the evolutionary relationship between *C. albicans* and *S. cerevisiae* Dnf1 and Dnf2 proteins from sequence comparisons alone. However, while *Ca*Dnf1 and *Ca*Dnf2 both localize to the plasma membrane, *Ca*Dnf2 is the primary flippase and is responsible for 70% of the PC uptake and 85% of the GlcCer uptake, with *Ca*Dnf1 catalyzing the remaining transport activity for these two substrates. These results are consistent with *S. cerevisiae* where Dnf2 is also the primary transporter of NBD-GlcCer and NBD-PC at the plasma membrane in comparison to Dnf1 (48). In addition, the transport activity and substrate preferences for ScDnf2 and CaDnf2 were nearly identical when both were expressed in *S. cerevisiae*. Based on these analyses, we decided to name C3_03250W_A and C5_00570W_A as *Ca*Dnf1 and *Ca*Dnf2 respectively.

In contrast, the sequence comparisons showed a clear evolutionary relationship between the *C. albicans* and *S. cerevisiae* P4-ATPase β-subunits (Cdc50, Lem3 and Crf1). Because of the high degree of sequence similarity between Lem3 protein from the two species (46% identity, 63% similarity), we initially thought that heterologously expressed *Ca*Dnf2 in *S.cerevisiae* would form a functional heterodimeric flippase with the *Sc*Lem3 in a *dnf1,2*Δ background. However, this was not the case and the cells expressing *Ca*Dnf2 alone failed to transport lipid substrate across the plasma membrane. Therefore, we co-expressed the *Ca*Lem3 and *Ca*Dnf2 together in the *Scdnf1,2*Δ strain and this yielded a functional transporter. It will be interesting to explore how the residues at the interface between the two subunits have changed over evolutionary time in a manner that prevented the cross-species interaction. Our data is consistent with the previous findings showing that the absence of β subunits affects trafficking of flippases to plasma membrane and flippase activity and confirm the identity of *Ca*Lem3 (20, 21, 58, 60).

*Ca*Dnf2 is a broad substrate specificity lipid transporter like *Sc*Dnf2 and transports GlcCer, PC and PE. We hypothesized that *Ca*Dnf2 utilizes a substrate translocation path like *Sc*Dnf2 based on the conserved residues critical for PC/GlcCer selection. Consistently, mutations in the predicted TM1 and TM4 substrate entry site perturbs the transport of PC and GlcCer. Substitution of the *Ca*Dnf2 TM1 substrate entry site GA residues with a QQ PS transport motif provides a gain-of-function PS transport activity, attenuates PC/GlcCer transport but does not influence PE transport. The affects of these mutations on substrate specificity are nearly identical in *Ca*Dnf2 and *Sc*Dnf2 (Fig.2,(21, 48, 60)). Thus, the mechanisms of substrate recognition and transport are well-conserved (58, 59).

*Ca*Dnf2 displays a polarized localization to the buds in yeast-form cells and to newly formed hyphal extensions during filamentous growth, whereas *Ca*Dnf1 is more uniformly distributed around both the mother and daughter cells. Consistent with this localization pattern, we find *Cadnf2Δ/*Δ cells display a defect in the yeast to hyphal transition and in the elongation of hyphae, leading to smaller colonies on Spider medium than the WT strain. In contrast, disruption of *CaDNF1* had no significant influence on polarized growth phenotypes. This is in contrast to *Saccharomyces cerevisiae*, where Dnf1, Dnf2 and the distantly related PS/PC flippase Dnf3 appear to act redundantly for many processes, including polarized growth underlying extension of the mating projection, or schmoo, and pseudohyphal growth (56, 72).

Mechanistically, the defects in cell polarity observed in *S.c.dnf1,2,3*Δ cells have been associated with perturbation of signaling molecule (protein and lipid kinases and Cdc42) interaction with the inner leaflet of the plasma membrane (72, 73). In the case of Cdc42, control of Dnf1 and Dnf2 over inner leaflet PS and PE were proposed to be critical for Cdc42 dynamics at the plasma membrane. In addition, the ability of Dnf3 to transport PS is thought to be important for its role in pseudohyphal growth (56). Our data, however, argue against a role for PS transport by *Ca*Dnf2 in hyphal growth because this flippase does not appear to transport PS. In addition, the *Ca*Dnf2 TM1 GA to QQ mutation allows PS transport and does not alter PE transport, yet this mutant fails to support hyphal growth.

Both the TM1 and TM4 mutations in *Ca*Dnf2 tested here strongly attenuate GlcCer transport. The inability of these *Ca*Dnf2 mutants to support hyphal growth, along with the known requirement for GlcCer in this process, makes it likely that the ability of *Ca*Dnf2 to transport GlcCer is critical for promoting polarized growth. We speculate that specific requirement for *Ca*Dnf2 in hyphal growth is due to both its polarized localization to the sites of growth and its stronger preference for GlcCer relative to *Ca*Dnf1. However, PC transport is also reduced by the TM1 and TM4 mutations used in this study and so it remains possible that PC is the critical *Ca*Dnf2 substrate needed to support polarized growth. Identification of better separation-of-function mutations in *Ca*Dnf2 are needed to clearly distinguish the roles of PC and GlcCer transport. Identifying the influence of these substrates, or their downstream metabolites, on signaling events important for polarized growth also requires further work.

P4-ATPases regulate vesicle budding in the endocytic and secretory pathways, and the interplay of these two pathways is critical for polarized growth. *Sc* Dnf1 and Dnf2 have been implicated in endocytosis at low temperatures and recycling of endocytosed protein back to the plasma membrane (50, 51). In the filamentous fungus *Aspergillus nidulans*, the Dnf1/2 ortholog DnfA localizes to the plasma membrane and the Spitzenkörper, which is center for hyphal growth and morphogenesis. The *dnfA*Δ mutant displays defects in hyphal growth and germination (44). Thus, the hyphal growth defects in mutants lacking *Ca*Dnf2 or flippase activity mutants could be due to defects in endocytosis or exocytic vesicle budding. It is also possible that *Ca dnf2* mutants downregulate the expression of hyphal growth specific genes important for virulence (43).

The *Ca dnf2*ΔΔ mutants also showed a reduced ability to infect mice as measured by fungal burden of the kidneys, a phenotype that was exacerbated in the *Ca dnf1*ΔΔ *dnf2*ΔΔ double mutant. Given the importance of hyphal growth to virulence, we suspect that the *Ca dnf2*ΔΔ mutant defects in polarized growth underlie this reduction in virulence. We also found that *Ca dnf2*ΔΔ mutants deficient in GlcCer/PC transport display the same defect in colonizing the kidneys of mice, correlating with the effects seen on hyphal growth. However, the more severe effect of the *Ca dnf1*ΔΔ *dnf2*ΔΔ double mutant on infection implies an additional role for *Ca*Dnf1 in pathogenesis, which may relate to its role in establishing membrane asymmetry. Alteration in plasma membrane asymmetry causes copper sensitivity in *Candida albicans* and host innate immune cells utilize copper to attack pathogens. Copper specifically binds to PS and PE and alters membrane permeability(74, 75). Deletion of *Candida albicans* flippases Dnf1, Neo1 or Drs2 causes sensitivity to copper, suggesting important roles for membrane asymmetry in protecting fungal pathogens from the host defense systems (76). Indeed, the virulence of several fungal pathogens is perturbed when P4-ATPases genes are disrupted, and it is possible that a general loss of membrane asymmetry makes the fungi more sensitive to the host defenses. Therefore, these P4-ATPases are attractive targets for antifungal drug development.

## Materials and methods

### Ethics statement

All animal work used in this study was done under an animal protocol (0016) that was previously approved by the University of Tennessee, Knoxville Institution Animal Care and Use Committee (IACUC) and was done in accordance with the National Institute of Health’s (NIH) ethical guidelines for animal research.

#### Phylogenetic tree analysis

Protein sequence of putative P4-ATPases were retrieved from Uniprot database. The accession number of the sequences used for analysis. *H. sapiens*: Q9Y2Q0 (HsAT8A1), Q9NTI2 (HsAT8A2), O43520 (HsAT8B1), P98198 (HsAT8B2), O60423 (HsAT8B3), Q8TF62 (HsAT8B4), O75110 (HsATP9A), O43861 (HsATP9B), O60312 (HsATP10A), O94823 (HsATP10B), Q9P241 (HsATP10D), P98196 (HsATP11A), Q9Y2G3 (HsATP11B), Q8NB49 (HsATP11C), Q9NV96(HsCdc50A), Q3MIR4(HsCdc50B); Saccharomyces cerevisiae: P40527 (ScNeo1p), P39524 (ScDrs2p), P32660 (ScDnf1p), Q12675 (ScDnf2p), Q12674 (ScDnf3p), P42838(ScLem3), P25656(ScCdc50), Q12674 (ScCrf1),; *Cryptococcus neoformans*: J9VZ19 (Apt1p), J9VQH2 (Apt2p), J9VGP8 (Apt3p), J9VM87 (Apt4p), J9VW44 (CnCdc50); *Candida albicans*: A0A1D8PDD2 (CaNeo1p), Q5ADR3 (CaDrs2p), A0A1D8PMY6 (CaDnf1p), A0A1D8PJN3 (CaDnf2p), A0A1D8PLQ7 (CaDnf3p), A0A1D8PHB8 (CaLem3), A0A1D8PQ78 (CaCdc50), CQ5AG97 (CaCrf1); *Aspergillus fumigatus*: Q4WCQ6 (AfDnfA), Q4X1T4 (AfDnfB), Q4WPR7 (AfDnfC), Q4WD94 (AfDnfD), A0A229W6V2 (AfCdc50). The sequence alignment was performed using MUSCLE and phylogenetic tree was created using Maximum Likelihood method in MEGA11 software.

#### Reagents

All yeast culture reagents were purchased from Sigma–Aldrich, Fisher scientific and Fisher bioreagents. NBD-lipids used in this study were purchased from Avanti Polar Lipids. *S.cerevisiae* strains were grown in YPD and SD selective media. *C*. *albicans* strains were grown in YPD media (1% yeast extract, 2% peptone, 2% dextrose) at 30°C while shaking at 250 rpm. YPM media (1% yeast extract, 2% peptone, 2% maltose) was used for flipping out the *SAT1*-flipper cassette.

#### Yeast strains and plasmid construction

Yeast strains and plasmids used in the study are listed in Table 1. The yeast strains were transformed using standard lithium acetate transformation. *Ca*Dnf1 and *Ca*Dnf2 were tagged on C-terminus with mNeonGreen with PCR-based gene tagging using pBR895 template (77). *Candida albicans* knockouts were generated using the SAT1 flipper strategy (78).

**Table 1:**
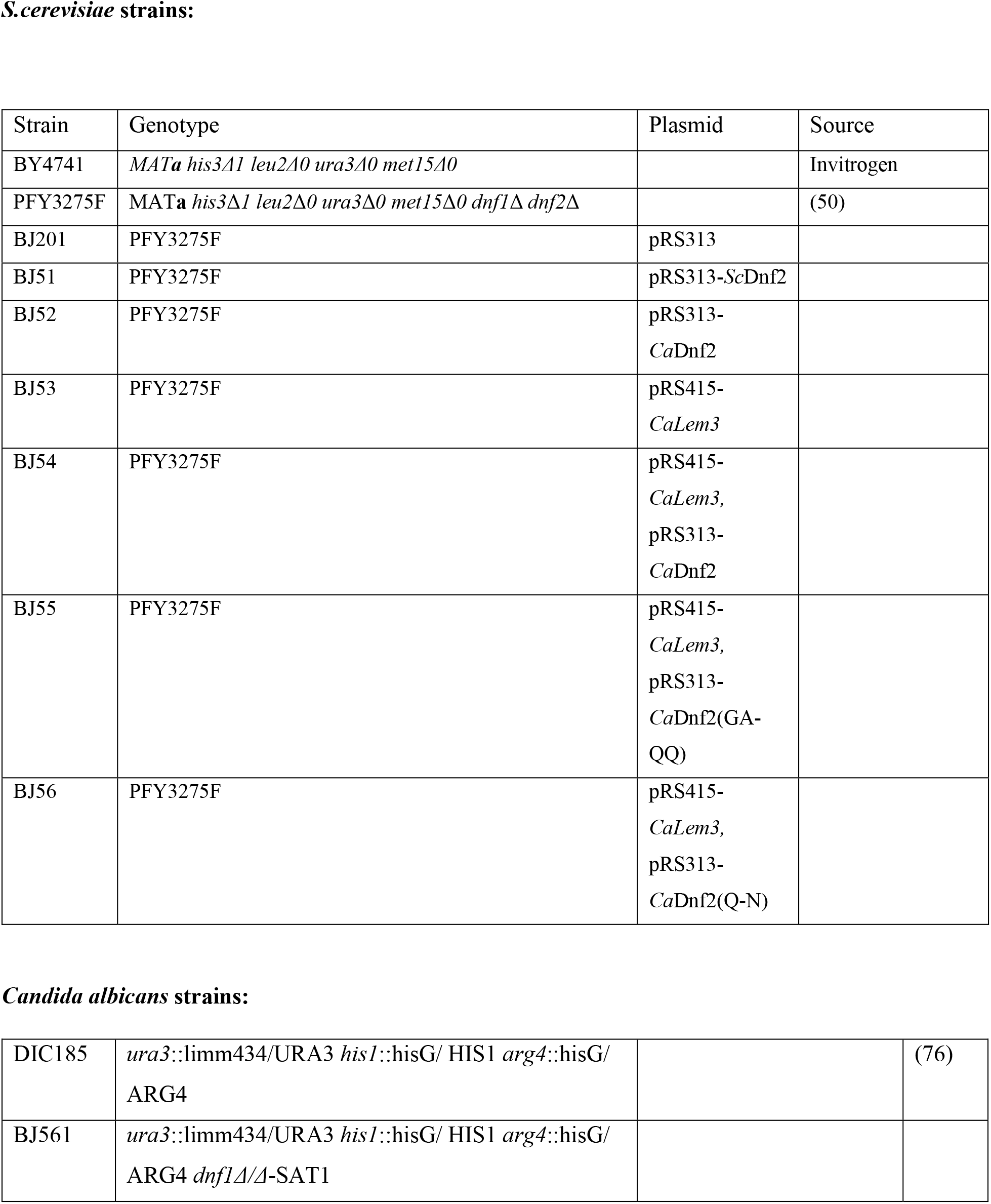
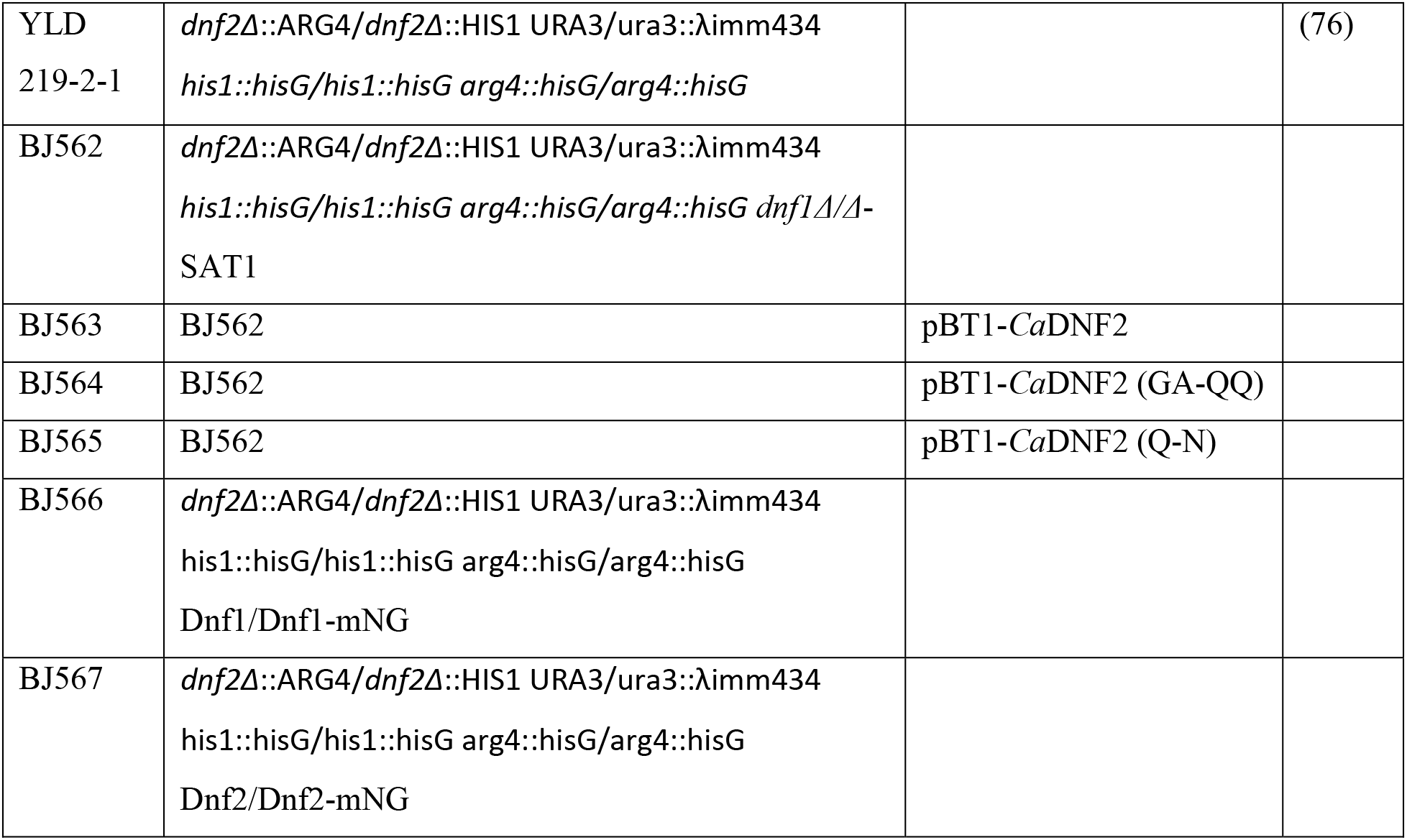
Strains and Plasmids used in the study.

| Strain | Genotype | Plasmid | Source |
| --- | --- | --- | --- |
| BY4741 | *MATa his3Δ1 leu2Δ0 ura3Δ0 met15Δ0* |  | Invitrogen |
| PFY3275F | *MATa his3Δ1 leu2Δ0 ura3Δ0 met15Δ0 dnf1Δ dnf2Δ* |  | (50) |
| BJ201 | PFY3275F | pRS313 |  |
| BJ51 | PFY3275F | pRS313-*ScDnf2* |  |
| BJ52 | PFY3275F | pRS313-<br>*CaDnf2* |  |
| BJ53 | PFY3275F | pRS415-<br>*CaLem3* |  |
| BJ54 | PFY3275F | pRS415-<br>*CaLem3*,<br>pRS313-<br>*CaDnf2* |  |
| BJ55 | PFY3275F | pRS415-<br>*CaLem3*,<br>pRS313-<br>*CaDnf2*(GA-QQ) |  |
| BJ56 | PFY3275F | pRS415-<br>*CaLem3*,<br>pRS313-<br>*CaDnf2*(Q-N) |  |
| DIC185 | *ura3::limm434/URA3 his1::hisG/ HIS1 arg4::hisG/ ARG4* |  | (76) |
| BJ561 | *ura3::limm434/URA3 his1::hisG/ HIS1 arg4::hisG/ ARG4 dnf1Δ/Δ-SAT1* |  |  |
| YLD<br>219-2-1 | *dnf2Δ::ARG4/dnf2Δ::HIS1 URA3/ura3::λimm434*<br>*his1::hisG/his1::hisG arg4::hisG/arg4::hisG* |  | (76) |
| BJ562 | *dnf2Δ::ARG4/dnf2Δ::HIS1 URA3/ura3::λimm434*<br>*his1::hisG/his1::hisG arg4::hisG/arg4::hisG dnf1Δ/Δ-*<br>SAT1 |  |  |
| BJ563 | BJ562 | pBT1-*Ca*DNF2 |  |
| BJ564 | BJ562 | pBT1-*Ca*DNF2 (GA-QQ) |  |
| BJ565 | BJ562 | pBT1-*Ca*DNF2 (Q-N) |  |
| BJ566 | *dnf2Δ::ARG4/dnf2Δ::HIS1 URA3/ura3::λimm434*<br>*his1::hisG/his1::hisG arg4::hisG/arg4::hisG*<br>Dnf1/Dnf1-mNG |  |  |
| BJ567 | *dnf2Δ::ARG4/dnf2Δ::HIS1 URA3/ura3::λimm434*<br>*his1::hisG/his1::hisG arg4::hisG/arg4::hisG*<br>Dnf2/Dnf2-mNG |  |  |

#### NBD-Lipid uptake assay

The yeast cells were grown to the mid-log phase, and 500 μl of culture were collected for each strain. NBD-PC and NBD-GlcCer were dried and resuspended in 100% ethanol and then added to the designated ice-cold growth medium at a final concentration of 2 μg/ml. Final ethanol volumes were ≤0.5%. The cells were incubated on ice with prechilled NBD lipid + medium for 30 min. Following incubation, the cells were washed twice with prechilled SA medium (SD medium + 2% (w/v) sorbitol + 20 mm NaN3) supplemented with 4% (w/v) fatty acid-free BSA. Finally, the cells were resuspended in SA medium with 5 μm propidium iodide (PI), and NBD-lipid uptake was immediately measured by flow cytometry. The order in which samples were prepared and processed was varied in each experiment to reduce potential experimental error. Each experiment consisted of three independent biological replicates for each strain and repeated three times.

### Flow cytometry

The flow cytometry experiments were performed on a three-laser BD LSRII (BD Biosciences) operating a FACS Diva 6.1.3 software package. Single-cell populations were identified by forward- and side-scatter as described previously (48, 49). Propidium iodide (PI) was used to exclude dead cell populations and those with lost membrane integrity, and the NBD signal was measured with FITC filters (530/30-nm band-pass filter with 525-nm long-pass filter). At least 10,000 cells were analyzed per experimental replicate.

#### Fluorescence microscopy

To visualize mNeonGreen tagged proteins, the cells were grown to mid-logarithmic phase in YPD medium. The cells were washed with fresh media three times and resuspended in fresh SD complete medium. The cells were mounted on glass slides and observed immediately at room temperature. The images were acquired using a DeltaVision Elite imaging system (GE Healthcare Life Sciences) equipped with a 100× objective lens followed by deconvolution using softWoRx software (GE Healthcare Life Sciences).

#### Hyphal growth and filamentation assay

Filamentation of *Candida albicans* cells was assessed using Spider medium (1% Nutrient broth, 1% mannitol and 0.2% K2HPO4). We then spread 10 to 50 cells in 100 μL of water onto plates and incubated them for 7 days at 37°C. The colonies were captured using a stereo microscope Nikon SMZ745T with 7.5X magnification. Filamentation in liquid media was assessed by inoculating cells into the filamentation media at an appropriate dilution and growing them at 37°C with shaking for 4 hrs. Cells were collected and mounted on a glass slide and observed under the microscope. Images were captured using the DeltaVision Elite imaging system. The filament length was measured using line tool of ImageJ 1.53k.

#### Mouse Model

Outbred ICR mice (ENVIGO) were used for all experiments in this study. To inoculate, 50 ml cultures of *C. albicans* strains were grown overnight in YPD media at 30°C while shaking at 225 rpm. The next day, cells were transferred to 50ml conical tubes (Thomas Scientific) and spun at 3,500 rpm for 5 minutes. Cells were subsequently washed 2 times with 25ml of PBS and then counted using a hemocytometer. Cells were then diluted to 1×10^7^ cells/ml in PBS. Mice were then intravenously injected via the lateral tail vein with 0.1ml of the prepared cell suspension. 5 days post infection, mice were euthanized, and their kidneys were harvested to assess fungal burden. Kidneys were placed in pre-weighed whirl-pack bags (Thermo Fisher Scientific) containing 1ml of water. The bags were weighed once more to determine the weight of the kidneys and then the tissues were homogenized. Serial dilutions of the tissue homogenates (10^−1^, 10^−2^, 10^−3^) were created, and 1ml of each dilution was added to 15ml of YPD and plated. Plates were then left to incubate for 2 days at 30°C to determine fungal colony forming units (CFU) per gram of tissue.

#### Statistical analysis

The statistical analyses were performed using GraphPad Prism 8.4.1. Variance was calculated using a one-way ANOVA test and comparisons with WT strains were tested with Tukey’s post hoc analysis. The p values represent the significance of the data: *, p < 0.05; **, p < 0.01; ***, p < 0.001; and ****, p < 0.0001.

## Data availability

The primary data will be available upon request. No material transfer agreements are required for data accessibility.

## Acknowledgments

We thanks Prof.James Konopka at Stony brook university for sharing the *C.albicans* strains and plasmids. The yeast flow cytometry experiments were performed in the Vanderbilt Medical Center (VMC) Flow Cytometry Shared Resource.

## Funding and additional information

This work was supported by National Institutes of Health Grants R35 GM144123-01 (to T. R. G.) and 1R01AI53599-01 (to T.B.R). The VUMC Flow Cytometry Shared Resource is supported by NIH grants to the Vanderbilt Ingram Cancer Center (P30-CA68485) and the Vanderbilt Digestive Disease Research Center (P30-DK058404). The content is solely the responsibility of the authors and does not necessarily represent the official views of the National Institutes of Health.

